# A Tomato Genome From The Italian Renaissance Provides Insights Into Columbian and Pre-Columbian Exchange Links And Domestication

**DOI:** 10.1101/2022.06.07.495125

**Authors:** Rutger Vos, Tinde van Andel, Barbara Gravendeel, Vasia Kakakiou, Ewout Michels, Elio Schijlen, Anastasia Stefanaki

## Abstract

The history of the tomato’s interactions with humans spans both a pre-Columbian period of millennia of cultivation and domestication in South and Central America as well as the plant’s rise to becoming one of the world’s major cash crops following the arrival of Europeans in the Neotropics and the subsequent Columbian Exchange. In the process, the influence of successive regimes of artificial selection gave rise to an entangled species complex comprised of a wild ancestor and numerous cultivars exhibiting a great diversity of fruit phenotypes (often sorted in ‘cherry’ and big types). Here, we provide a snapshot into these dynamics by presenting the ancient nuclear and plastid genomes of one of the oldest tomatoes still in existence: the almost half a millennium old *En Tibi* specimen from the Italian Renaissance. By placing the genome skimming data obtained from this specimen within the phylogenomic context of wild relatives and Neotropical tomato cultivars and land races, we are able to show that the most likely geographic origins of the specimen’s immediate ancestors, i.e. whence the exchange of this particular lineage originated, are around the Gulf of Mexico. We also show that this specimen was less inbred than present-day tomatoes, shedding light on the genomic signatures of historical domestication processes. Probing deeper into the ancestry that shaped extant genetic diversity within the complex, we reconstruct multiple ancestral populations with diffuse but distinct signatures in cherry and big tomatoes. The geographic structure of these ancestries within the nearest wild relative of tomatoes and the differential extent to which these ancestries are represented in domesticates points to early cultivation and human-assisted dispersal of tomatoes originating from the northern extreme of the natural range of the species. Our findings illustrate the inferences that can be drawn from ancient DNA extracted from herbarium specimens and historic plant collections. At the same time, we stress that our findings draw on multiple disciplines including ethnobotanical and historical research and on the (agri)cultural contributions of a variety of world cultures.

## Introduction

The fruits of the domesticated tomato plant (*Solanum lycopersicum* L., Solanaceae) are an important food crop, globally ranking twelfth by tonnage (1) with 244 million metric tonnes produced across 176 countries in 2018 (2). Although originating in South America, yields – mostly in greenhouses – are currently highest in the Netherlands, with a national average of about 509 tonnes per hectare (2). The domesticated tomato belongs to the section *Lycopersicon* of the large genus of the nightshades, *Solanum*, a genus that also contains the economically important potato (*Solanum tuberosum* L.) and eggplant (aubergine, *Solanum melongena* L.) among many other species. The section *Lycopersicon* contains a well-supported clade (3) of four orange- or red-fruited tomato species: *S. lycopersicum* L., *S. pimpinellifolium* L., *S. cheesmaniae* (Riley) Fosberg, and *S. galapagense* S. C. Darwin & Peralta. The latter two species are only found on the Galapagos Islands, while *S. pimpinellifolium* L. occurs in the wild mainly in South America, especially in Chile, Peru and Ecuador. The domesticated *S. lycopersicum* L. can be found in tropical regions across the globe due to the feralisation of locally cultivated plants (4).

The domesticated tomato is commonly subdivided further into the cherry tomato *S. lycopersicum* var. *cerasiforme* and *S. lycopersicum* var. *lycopersicum*, which is the large-fruited tomato. (For brevity’s sake, we will occasionally refer in this manuscript to *S. pimpinellifolium* L. as “P” and the domesticated varieties as “C” and “B”, for “cherry” and “big”, respectively). A common model holds that domestication of tomatoes was a two-step process initially resulting in C from P, followed by further improvement leading to B (5). A more recent study proposes, instead, that C initially evolved naturally (6). Comparison of regions with reduced nucleotide diversity between P/C on one hand and C/B on the other reveals that different sets of genes were affected in two distinct stages of evolutionary change (5), findings that are broadly compatible with either model. However, analysis of population structure shows that multiple ancestral populations and introgression have shaped C’s diffuse placement in relation to both B and P such that a progression of fully sorted, nested monophyly of the form P→C→B does not occur; instead, a more complex pattern of interdigitation among accessions generally results (3, 7, 8). Nevertheless, broad geographic trends in the stages of tomato domestication in the pre-Columbian era can be discerned.

The initial stages of the evolutionary process occurred in the Andes (present-day Peru and Ecuador), after which additional stages took place in Mesoamerica, perhaps much later, as inferred from diminishing genetic diversity in C from Ecuador to Mexico (5, 7, 9). Such reductions in genetic diversity are a common feature of domestication in general. They have been demonstrated at a broader and more exacerbated scale among numerous accessions of landraces and cultivars in the tomato clade (3). It remains unclear how and where the second stage of pre-Columbian tomato domestication occurred. The genetic evidence suggests two possibilities; either tomatoes were transported and improved gradually from Ecuador to Colombia and Costa Rica to Mesoamerica, or there was direct transport of domesticated tomatoes to Mesoamerica, where further improvement took place in its entirety (7).

Domesticated, large-fruited (B) tomatoes were introduced to Europe during the first half of the sixteenth century as part of the Columbian Exchange (10). Not long after their first introduction, tomatoes were grown as exotic curiosities in gardens of the elite, while naturalists of the time started describing them and collecting them (4, 11, 12). The Italian physician Pietro Andrea Mattioli (1501-1578) wrote the first botanical description of tomatoes in 1544 in his *Commentaries on Dioscorides*. After describing the eggplant, he writes: ‘Another species has been brought to Italy in our time, flattened like red apples and composed of segments, green at first and of a golden colour when ripe, and they are also eaten in the same way’ [as eggplants] (11, 13, 14). In a later edition of this book, he named these plants *pomi d’oro* (golden apples) (15).

Sixteenth-century naturalists did not just describe tomatoes but also collected them in herbaria. Several of these early tomato specimens have survived to this day in historic herbaria, most of which of Italian origin (for an overview of sixteenth-century tomato specimens see (14)). One of the oldest herbaria containing a tomato specimen is preserved in the collection of Naturalis Biodiversity Center (16). It is bound in leather and gold and bears the inscription *En Tibi Perpetuis Ridentem Floribus Hortum* (‘Here for you, a smiling garden of everlasting flowers’). This inscription and the luxurious features of the book imply that it was prepared to be offered as a present to a person in power, presumably the Habsburg emperor Ferdinand I (1503-1564) (17). The *En Tibi* herbarium was made in the area of Bologna around 1558 (17, 18). It contains 473 plant specimens across 455 taxa and 97 different families (17). Most of these plants are native to northern and central Italy (17). Still, the herbarium also contains several taxa native to Asia and five species native to South America, among which is one of the oldest extant tomato specimens. The plant material and species list for the herbarium were most likely provided by the Italian physician Francesco Petrollini, who was also the maker of one of the largest and oldest herbaria preserved today, the so-called Erbario B (“Erbario Cibo”) kept in the Biblioteca Angelica in Rome (18).

However, Petrollini was probably not involved in the actual construction of the herbarium considering the numerous clerical errors observed in the plant names (Fig. 1). DNA evidence of human hairs found glued underneath several plants on the herbarium sheets revealed that at least four individuals were involved in the making of this herbarium (18).

**Figure 1 –.**
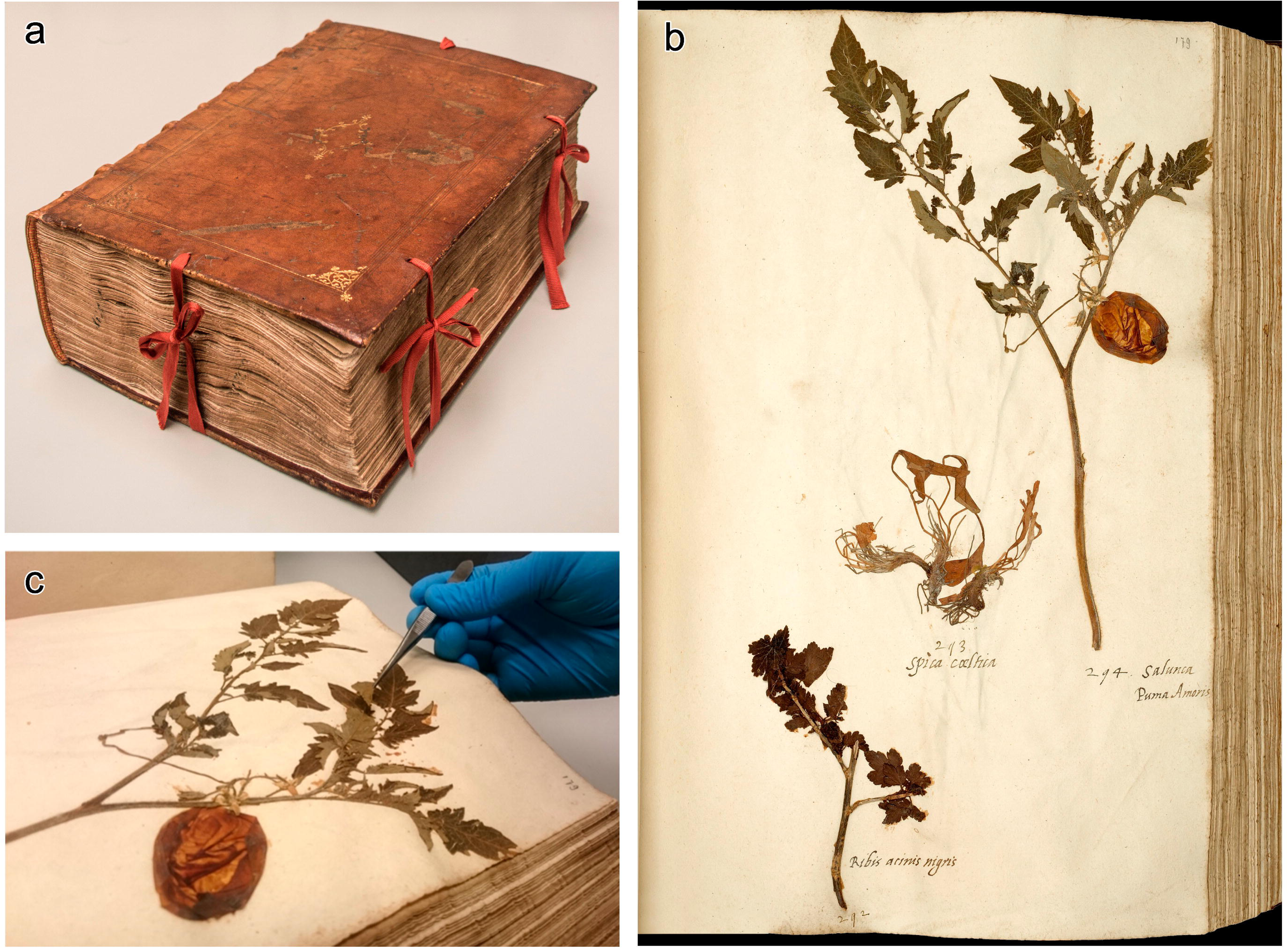
The *En Tibi* specimen. a) The book herbarium *En Tibi Perpetuis Ridentem Floribus Hortum* (‘Here for you, a smiling garden of everlasting flowers’), compiled ca. 1558. b) The page containing the tomato specimen, top-right, labelled *Puma Amoris* (‘Love apple’) in context with other specimens on the same page. The name ‘Salunca’ was erroneously given to the tomato specimen by the scribe who wrote the plant names on the book, it actually refers to the previous specimen. c) Sampling of leaf material for ancient DNA extraction.

Modern DNA techniques can further elucidate the origins and characteristics of historic herbarium specimens, as exemplified in this study by the ca. 500-year-old *En Tibi* tomato specimen. Tomatoes are diploid and have a – for plants – relatively small genome of approximately 823.1 megabases (Mb) mainly comprised of low-copy DNA (19, 20). Due to this, a high-quality assembly of the Heinz 1706 cultivar with annotations locating approximately 30-35 thousand genes (20) has been available since 2012, which has spurred on several large-scale resequencing projects (3, 5). These projects have given researchers access to panels of hundreds of tomato cultivars, landraces and wild relatives mapped against a common reference.

The recent tomato resequencing studies have relied chiefly on germplasm collections in gene banks to obtain samples for sequencing. These, however, are not the only tomato accessions being curated. Natural history collections, such as herbariums, provide valuable additional information on phylogenomics, geographic structure and genetic variation. For example, a recent study demonstrated how phylogenomics of museum specimens could reconstruct the geographic origin of poorly annotated specimens using a panel of conspecifics with good georeferencing (21). Older specimens may yield especially exciting snapshots of genetic variation through space and time. For domesticated species, such snapshots are of distinct interest in that the dynamics that gave rise to their reduced genetic diversity are under debate. The conventional view (*sensu* Allaby et al. (22)) is that reduced genetic diversity results from an initial population bottleneck at the ‘domestication event’, following which there may even be an increase in diversity subsequent to agricultural expansion. Alternatively, genetic erosion may compound gradually due to the effects of ongoing artificial selection and breeding. Using archeogenomics, Allaby et al. demonstrated that the alternative model is best supported by the evidence in sorghum, maise and barley; i.e., there is hardly a discernible reduction in genetic diversity among ancient specimens of these domesticated crops at all.

In light of the expanding resequencing data sets, progressing insights in the dynamics of domestication, and methodological advances in ancient DNA and bioinformatics, compelling questions surrounding the *En Tibi* specimen can be addressed in more detail than previously possible. This study interrogates the specimen’s ancient DNA to place the snapshot it represents within the broader context of tomato diversity. The first question we seek to address concerns the geographic origins of the variety from which the specimen derives. The second question is that of genetic diversity through time under domestication and cultivation. From its historical provenance in Italy, we know that the specimen was cultivated and therefore presumably a domesticated tomato; however, it predates industrial selective breeding of tomatoes by centuries and should therefore provide some clues in the dynamics of genomic erosion over time. The final question is that of the cultivation and trade of pre-Columbian tomatoes. The specimen originates from the time when the tomato was just on the cusp of transitioning from a neotropical, indigenous heirloom to becoming a cosmopolitan cash crop. Presenting its genome within the context of the broader diversity of neotropical landraces and cultivars draws attention to the invaluable contributions of Amerindian agriculture in feeding the world.

## Materials and Methods

### Sampling and aDNA extraction

Ancient DNA was extracted from the *En Tibi* tomato specimen in the dedicated Ancient DNA facility of Naturalis Biodiversity Center by grinding a small leaflet fragment (<2 cm^2^, 4 mg, see Fig. 1C) with a Retsch Mixer mill (MM 400) using glass beads (0.5 mm diameter) for 5 min at 30 Hz. DNA extraction was performed following a modified CTAB-protocol (23). For digestion, a lysis buffer containing 50 mM Tris-HCl (Thermo Fischer Scientific), 3% 2-mercaptoethanol (Sigma Aldrich), and 0.25 mg/ml Proteinase K (Promega) was applied to the leaf powder. Strict cleaning regimes and clean lab practices (24, 25) were applied in all steps.

### DNA sequencing

Because a DNA quality check on a Bioanalyzer High Sensitivity DNA assay showed the *En Tibi* DNA was severely degraded, the material was first subjected to a damage repair procedure using NEBnext FFPE DNA repair mix. Subsequently, DNA was used for fragment end repair, A-tailing, barcoded adapter ligation and library amplification according to NEXTFLEX ChIP-Seq Kit guidelines (BIOO Scientific). Finally, the amplified library was double purified with Ampure XP beads (Beckman Coulter) quantified by Qubit fluorometric assay (Invitrogen), and library size was determined on a Bioanalyzer High Sensitivity DNA assay. The *En Tibi* library showing a peak size at 308 bp; 2.6 ng/uL concentration was further diluted and 12 pM was used for clustering on one lane of a single illumina Paired End flowcell using a cBot system (illumina). Sequencing was done on an illumina HiSeq2000 instrument using 2*125 nt Paired-End sequencing performed by the genomics facility of Wageningen University and Research. De-multiplexing of resulting data was carried out using Casava 1.8 software (illumina).

### Genome assembly and homologous data set creation

We removed any remaining forward and reverse adaptors using cutadapt (26) from the FASTQ data resulting from DNA sequencing. We then used fastp (27) to trim low-quality ends and perform an initial quality assessment on the results. As a reference genome, we used assembly release 2.50 of the genome of *S. lycopersicum* cv. Heinz 1706 (RefSeq GCF_000188115.3) and mapped our reads against this using minimap2 (28) with the optimisations for genomic short-read mapping (‘sr’). We then used the samtools (29) chain to add mate coordinates, insert size fields, and remove secondary and unmapped reads (‘fixmate’); to sort the mapped reads (‘sort’); and to remove duplicates (‘markdup’). For variant calling, we used the bcftools (30) chain to compute a pileup (‘mpileup’), call variants (‘call’), and remove (‘filter’) low mapping phred scores (QUAL<20). To assess the extent to which the ancient DNA had been affected by post-mortem sequence modification (31), we performed a mapDamage (32) analysis on the assembly.

To make unbiased comparisons between the *En Tibi* nuclear genome, which we found to be very fragmented and with shallow coverage, and other accessions, we located all autosomal regions with ≥10x coverage in the mapped assembly. We then selected all wild tomatoes, domesticated landraces and ‘Latin American cultivars’ (*sensu* Lin et al. (5)) from South and Central America that were resequenced in the 360-tomato project (resulting in a panel of 114 genomes). From these, we extracted all variants that occur within those same ≥10x coverage regions of the *En Tibi*. We homologised the resulting variants by importing them into a relational (SQLite) database and traversing the chromosomes and their respective variant loci in numerically sorted order, resulting in a character/state matrix where each column comprises a genomic site where at least one accession has a variant.

Because the 360-tomato project does not include mapped plastid genomes, we also assembled the plastomes for the panel of 114 accessions using their raw short read submissions at SRA. For this, we used the same pipeline as we applied to the nuclear data (up to but not including the variant calling procedure) using the tomato reference plastome sequence NC_007898. As plastids are much more numerous than nuclei, especially in leaf material like we sampled from the *En Tibi* specimen, coverage was very high in all specimens, including the *En Tibi*. Out of the mapped assemblies, we extracted the consensus sequences for the 12 cpDNA barcode markers previously established (33) as most suitable for evaluating plant phylogeny at low taxonomic levels. This procedure was performed by computing an mpileup with bcftools of the complete plastome sequence for each accession, converting this to FASTA with PGDSpider (34), splicing out the marker locations (Table 1) and multiple sequence aligning them separately with MUSCLE (35), then concatenating the resulting MSAs with phyutility (36).

**Table 1:** Genomic coordinates of highly variable chloroplast markers. Markers were selected based on a previously published reference set. Genomic coordinates are with reference to plastome sequence NC_007898.

| Chloroplast marker | Genomic coordinates |
| --- | --- |
| *trnH-psbA* | 78-536 |
| *trnK* | 1,813-4,398 |
| *rps16-trnQ* | 6,065-7,144 |
| *rpoB-trnC* | 27,099-28,420 |
| *trnS(UGA)-trnG(GCC)* | 36,347-37,169 |
| *ndhC-trnV* | 51,750-52,845 |
| *rbcl-accD* | 58,117-58,875 |
| *petA-psbJ* | 64,857-65,924 |
| *psbE-petL* | 66,692-67,688 |
| *ndhF* | 111,505-113,718 |
| *rpl32-trnL* | 114,669-115,578 |
| *ycf1* | 125,297-130,972 |

### Dimensionality reduction and clustering

To elucidate the placement of the *En Tibi* specimen within the diversity of neotropical wild and domesticated tomatoes, we first performed dimensionality reduction and visual clustering analyses of the broad panel of 114 genomes. This panel includes the wild species *S. cheesmaniae* (Riley) Fosberg (n=3), *S. chilense* Dunal (n=1), *S. galapagense* S. C. Darwin & Peralta (n=1), *S. habrochaites* S. Knapp & D.M. Spooner (n=1), *S. neorickii* D.M. Spooner, G.J. Anderson & R.K. Jansen (n=1), *S. peruvianum* L. (n=3) in addition to numerous accessions of P (n=45), C (n=42), and B (n=17). The dimensionality reduction method we applied was t-SNE (37, 38), parameterised with a perplexity value of 25 (39). Visual inspection of the results (Fig. 2) showed that the *En Tibi* specimen is a domesticated tomato, which the analysis placed among C and B accessions, although without a clear tendency towards one type or the other under this low resolution. We then zoomed in further on the C/B/P complex, omitting the other wild species in a NeighborNet (40) analysis implemented in SplitsTree (41).

**Figure 2 –.**
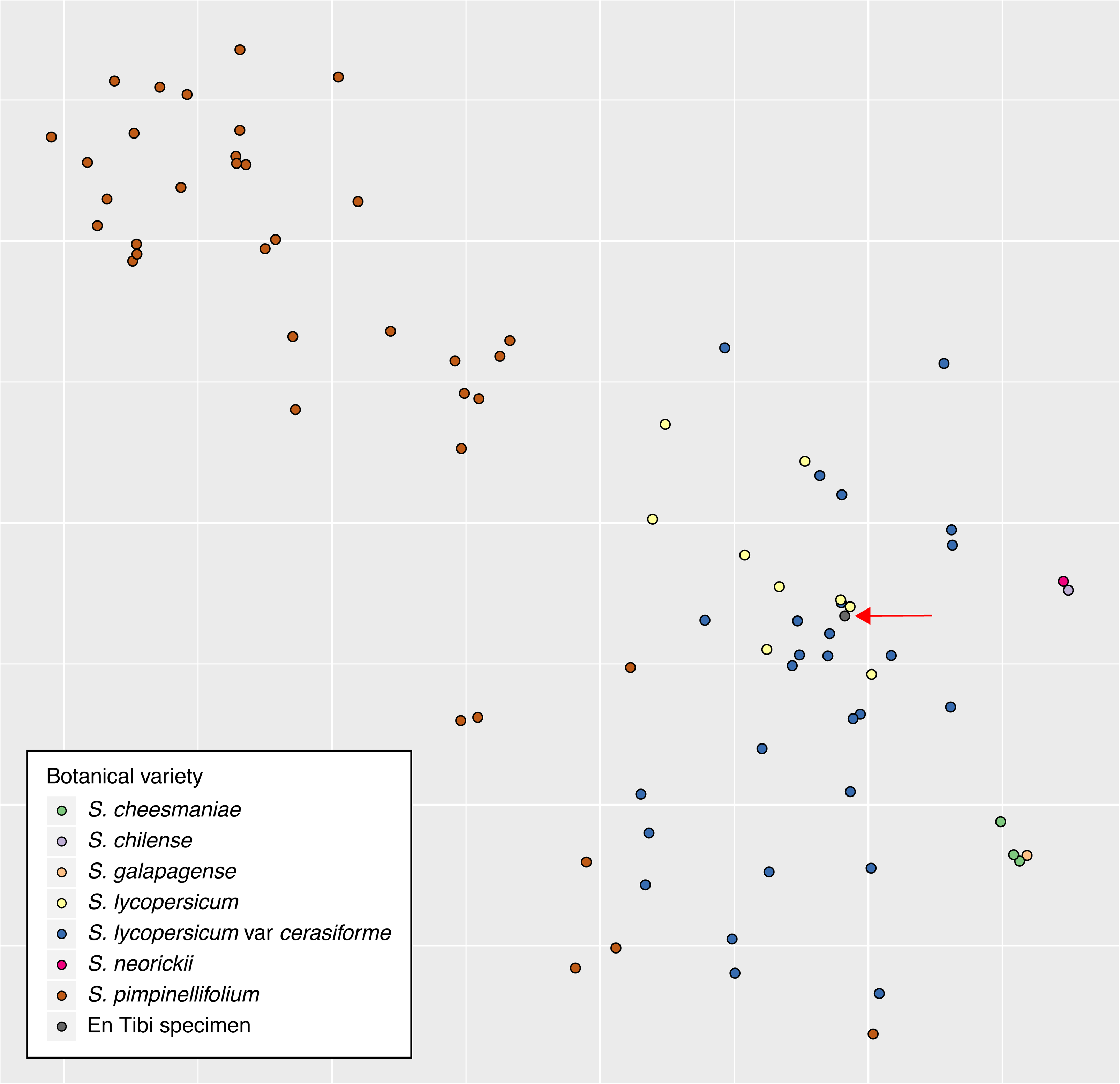
Dimensionality reduction of SNP results using t-distributed stochastic neighbor embedding (t-SNE). The variants data set consists of those encountered among the members of the panel, within genomic coordinates where the *En Tibi* specimen has mapping coverage >=10x.

Given the uniparental inheritance of the plastid markers, we expected a less ambiguous and less multidimensional signal in this dataset. We, therefore, omitted the t-SNE analysis on the plastid data and proceeded directly to network construction (42). As the number of variant sites in this data is much smaller than in the nuclear set, and due to its inheritance, a haplotype network approach was both feasible and appropriate. To this end, we constructed a Median-Joining network (43) as implemented in PopART 1.7 (44).

### Structure and heterozygosity

As previous authors have noted, the C/B/P complex shows admixture among the types resulting from introgression from multiple ancestral populations (8). This raises the question of the influences of these source populations on the *En Tibi* specimen. To assess this, we performed structure analyses on the character/state matrix using fastSTRUCTURE (45) across a range of numbers of hypothesised ancestral populations to derive the maximum likelihood estimate. To relate the hypothetical ancestral populations to geographic origins, we placed the fastSTRUCTURE results for the accessions of *S. pimpinellifolium* on the map of South America.

The complex that is the focus of our study includes both a wild species (P) and two domesticated forms (B and C), whose pre-Columbian domestication and breeding history spans a geographic extent that suggests repeated, successive founder events. As such, the genetic diversity in terms of heterozygosity likely spans a wide range, as has been demonstrated by previous authors (5, 7, 9). (The distribution of edge lengths across the NeighborNet in Fig. 3 also suggests this in qualitative terms). Within this distribution, the placement of the *En Tibi* specimen is of interest as a snapshot of the temporal and spatial dynamics that shaped genetic diversity during domestication and plant breeding. We, therefore, calculated heterozygosity for all accessions in the panel within the regions of sufficient (≥10x) coverage of the *En Tibi* and partitioned this by region (Andes or wider Neotropics), taxon/form (P, B, or C) and botanical status (wild, hybrid, cultivar, or landrace).

**Figure 3 –.**
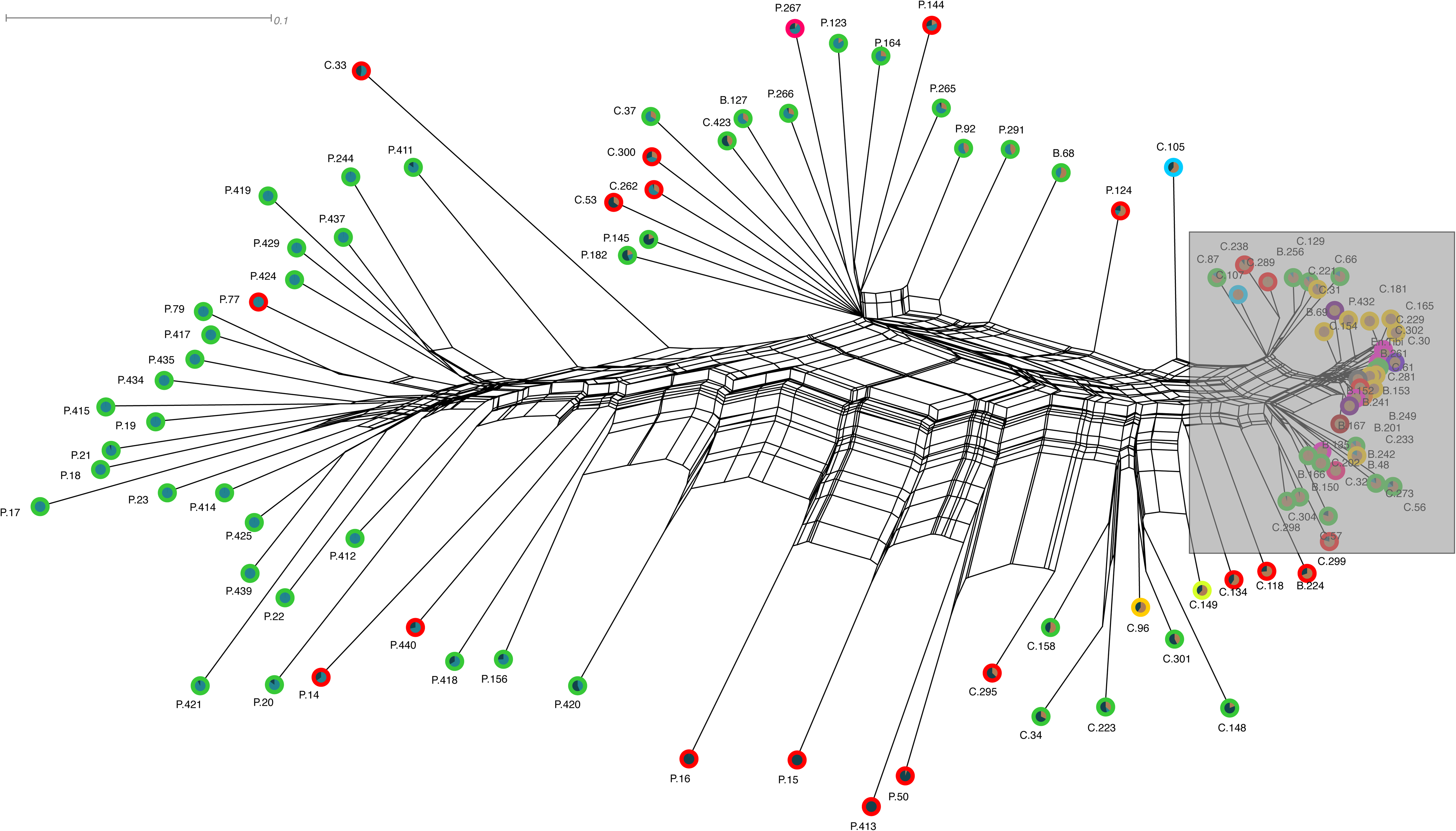
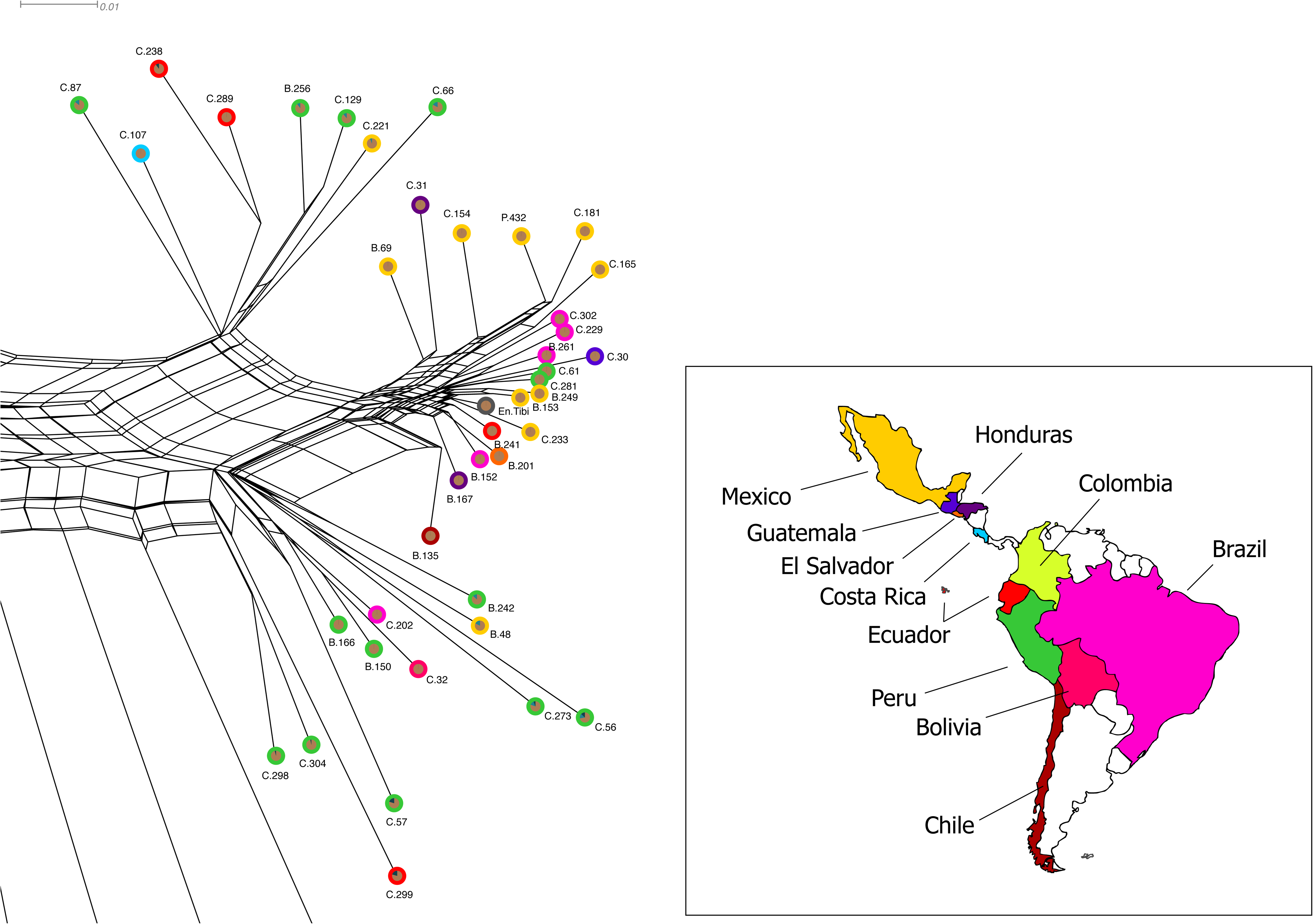
NeighborNet and fastSTRUCTURE results for the nuclear genome data, overview (a) and zoomed in on the *En Tibi* specimen (b). In these graphs, each terminal node represents an accession from Lin et al. (5) among which the *En Tibi* specimen is placed. The outer circle’s colour on each node corresponds with the collection country in the inset map in (b). Pie charts represent ancestry components detailed further in Figure 5.

## Results

### DNA sequencing and genome assembly

Sequencing yielded ±494 million PE reads (combined forward and reverse) containing ±60 gigabases with a GC content of ±60%. Base-calling quality distribution was ordinary, with 96.7% of bases called with a phred score greater than or equal to Q20 and 93.6% greater than or equal to Q30. After quality filtering, 97.5% of reads were retained. The expected read length on this sequencing platform is 125bp, which was the observed average; nevertheless, shorter reads were encountered at subjectively higher numbers than in fresh material, potentially indicating fragmentation of the source genome. Additional raw yield statistics are contained in the fastp and FastQC reports in the assembly archive of our supplementary materials deposition; the raw reads themselves are available from the NCBI SRA under BioProject PRJNA566320.

The mapping assembly of the nuclear genome yielded an alignment that we make available as a BAM file in our supplementary materials. We screened this BAM file using mapDamage, which indicated evidence of modest C → T misincorporations on the 5’ end and G → A misincorporations on the 3’ end of our reads. More detailed output results of the analysis are available in the mapdamage archive in the supplementary materials. Due to the deterioration of the source material and the need to minimise destructive sampling on this culturally and historically significant specimen, coverage across the nuclear genome was low: out of the approximately 823.1 million bases in the nuclear reference genome, only 9.9 million were covered at a depth greater than or equal to 10x, i.e. about 0.75% of the reference had a coverage we deemed sufficient for further analysis, requiring us to develop a highly fragmented genome skimming approach. To this end, we make available a relational database containing the 10x covered SNPs from the *En Tibi* specimen and all the well-covered SNPs from the selected accessions of the 360-tomato project that lie within the same genomic coordinates. This database is part of the snpdb archive in our supplementary materials.

Mapping the *En Tibi* reads to the plastid reference genome yielded good coverage: the reference was fully covered, with an average depth of 176x. Likewise, the selected accessions of the 360-tomato genome project also resulted in well-covered alignments. The average depth across these assemblies amounted to 172x. All plastid assemblies are available in the cp_dna archive in our supplementary materials.

### Dimensionality reduction and clustering

The results of the t-SNE analysis, available in the tsne archive of our supplementary materials, are shown in Figure 2. Most of the visualisation space is used to map the genetic diversity within the C/B/P complex. The other species are orientated in patterns that reflect their geographic and phylogenetic affinities: *S. cheesmaniae* and *S. galapagense*, both from the Galapagos Islands, are near each other, as are the Andean species *S. neorickii* and *S. chilense*. However, at this level of resolution, the analysis merely demonstrates that the *En Tibi* specimen is indeed a domesticated tomato, albeit qualitatively equidistant to both C and B accessions.

The results of the NeighborNet analysis on the nuclear SNPs (available in the neighbornet archive of the supplementary materials) are shown in Figure 3. The overall pattern (Fig. 3a) shows a concentration of P accessions on the left-hand side of the network. Towards the centre, there is increasing interdigitation of C accessions. The dense, short-branched cluster on the right-hand side, shown zoomed-in (Fig. 3b), contains C and B accessions. The accessions nearest to the *En Tibi* specimen were collected in Mexico, the closest two being B accessions and a more distance C accession. The median-joining analysis of the plastid markers resulted in the haplotype network shown in Figure 4. As in the t-SNE result, the broad patterns include a subgraph of Andean accessions (*S. neorickii* and *S. chilense*, here augmented with *S. peruvianum*) and a subgraph of accessions from the Galapagos Islands (*S. cheesmaniae* and *S. galapagense*). In this analysis, a more collapsed pattern of two large clusters emerges: one of South American C/P accessions around a central node and another of Mesoamerican C/B/P nodes, including the *En Tibi* specimen.

**Figure 4 –.**
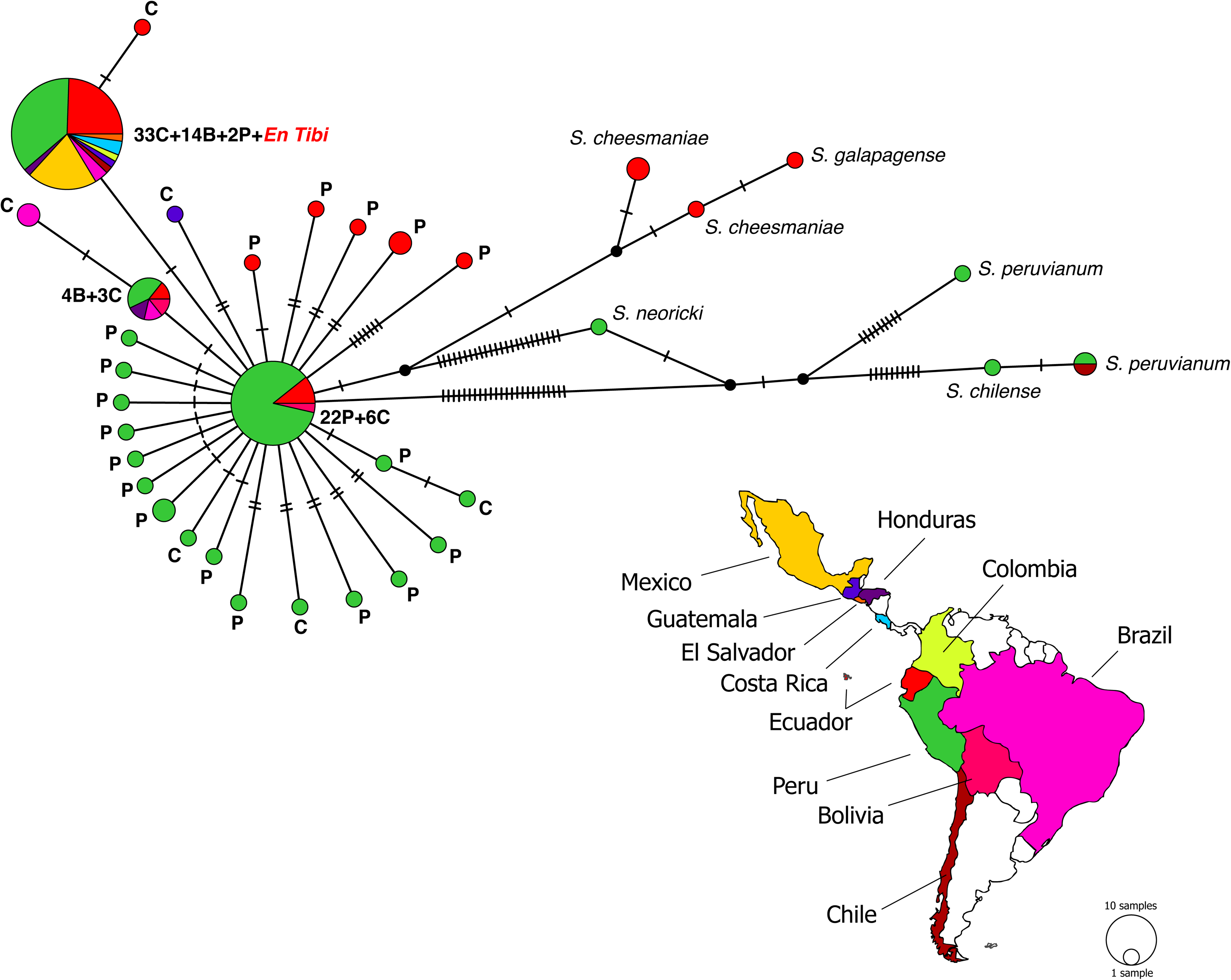
Median-joining network of the chloroplast data. Colours of the nodes correspond with the collection country in the inset map. Colours of the labels correspond with the species and varieties. Nodes containing multiple varieties are indicated with asterisks, and their respective compositions are shown in the figure legend.

Interestingly, a third cluster occurs in our data, consisting of C and B accessions spread out across South (Peru, Ecuador, Bolivia and Brazil) and Central (Honduras) America with an additional C haplotype budding off in Brazil. Further research would be needed to understand the historical processes that led to the development of this cluster.

### Structure and heterozygosity

The fastSTRUCTURE analysis resulted in a maximum likelihood estimate of three ancestral populations and assignments to these for the accessions in our data set, as shown in Figure 5a. Among C and P is an ancestral population practically absent from B. The *En Tibi* specimen is fully assigned to an ancestral population to which several accessions from both C and B were also fully assigned. In contrast, none of our P accessions are reconstructed to originate entirely from this population. The reconstructed ancestral populations show geographic structure within P. To illustrate this, we placed the P accessions and their reconstructed ancestry components on the map of the South American west coast, as shown in Figure 5b. The ancestral population to which many P accessions (but no C or B accessions) are fully assigned is concentrated in the South, in Peru. To the North, in the interior of Peru and Southern Ecuador, is a population to which some P accessions are fully assigned that also dominates the ancestry of some C accessions. Intermingled with that and with occurrences to the West is another reconstructed population, to which some C and some B accessions (including the *En Tibi* specimen) are fully assigned. The P accessions that have this ancestry the most were collected south of the Gulf of Guayaquil. The results of the fastSTRUCTURE analysis are available from the structure archive of the supplementary materials.

**Figure 5 –.**
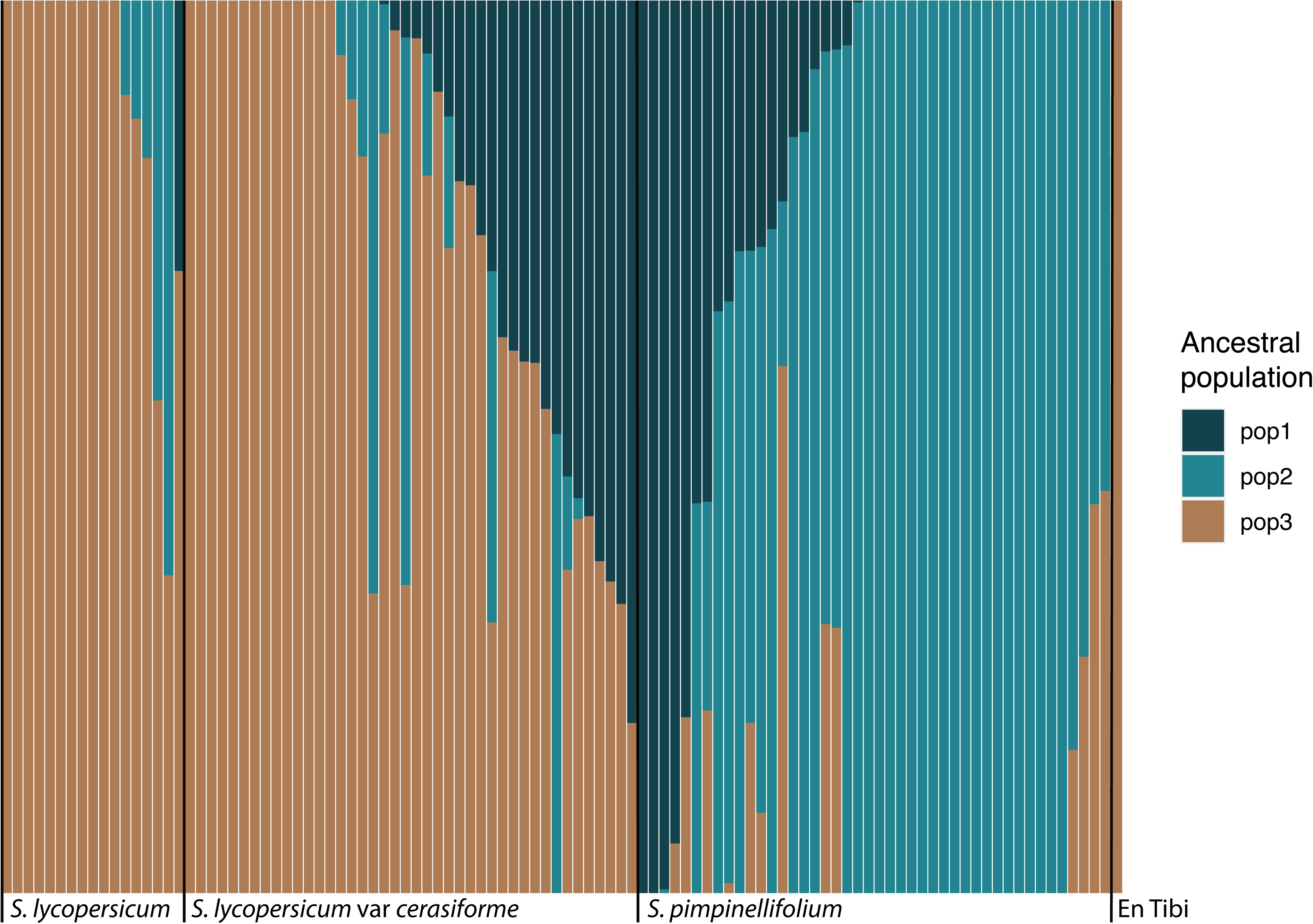
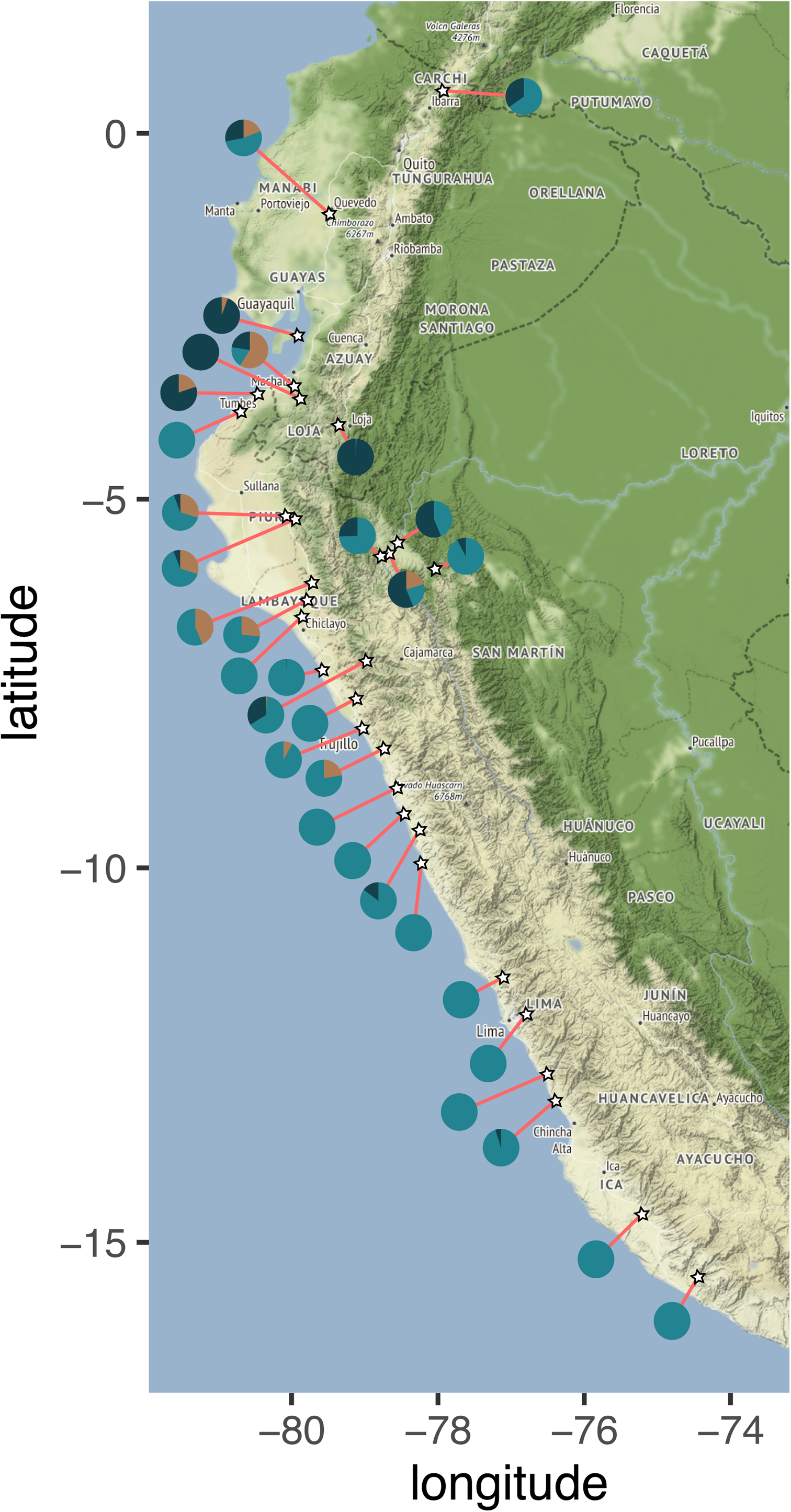
fastSTRUCTURE results. The analysis obtained a maximum likelihood estimate of three ancestral populations. Ancestry components for all accessions in the P/C/B complex are shown in (a). The geographic structure of the ancestry components inferred for *Solanum pimpinellifolium* L. are shown in (b).

In this study, we define heterozygosity as the ratio of the number of heterozygous SNPs over the number of homozygous SNPs within the genomic regions for which the *En Tibi* specimen had 10x coverage. In these regions, some SNPs might be so minimally covered (i.e. close or equal to 10x) that robust support for their heterozygosity could be questionable. We, therefore, consider both a ‘face value’ estimate of heterozygosity as well as a more conservative estimate, where the latter discards all SNPs where the coverage of the alternative allele is lower than 4x. In addition, we verified by eye that none of the heterozygous SNPs of the *En Tibi* specimen that we include were on trailing ends of our mapped reads to avoid including bases resulting from misincorporation. Due to these caveats, we, therefore, consider an interval of heterozygosity for this specimen and compare that with the more robustly supported point estimates from the other accessions.

The results of the heterozygosity analysis are shown in Figure 6, where the shaded area represents the interval for the *En Tibi* specimen. The results are also available from the heterozygosity archive in our supplementary materials. Figure 6a shows that heterozygosity is greatest in P, lower in C, and even lower in B. The *En Tibi* specimen’s estimated interval intersects with that of the heterozygosity seen in P. Figure 6b shows that heterozygosity is greatest in ‘wild’ tomatoes (*sensu* Lin et al. (5), who include feral specimens in this definition), less so in hybrids between ‘wild’ and cultivated tomatoes, and even lesser in cultivated tomatoes. In this pattern, the interval for the *En Tibi* intersects with that of the heterozygosity commonly seen in ‘wild’ tomatoes. In our data partitioning schemes, heterozygosity is greatest in the Andean region and lower in the wider Neotropics. The results of these analyses are available in the heterozygosity archive of our supplementary materials.

**Figure 6 –.**
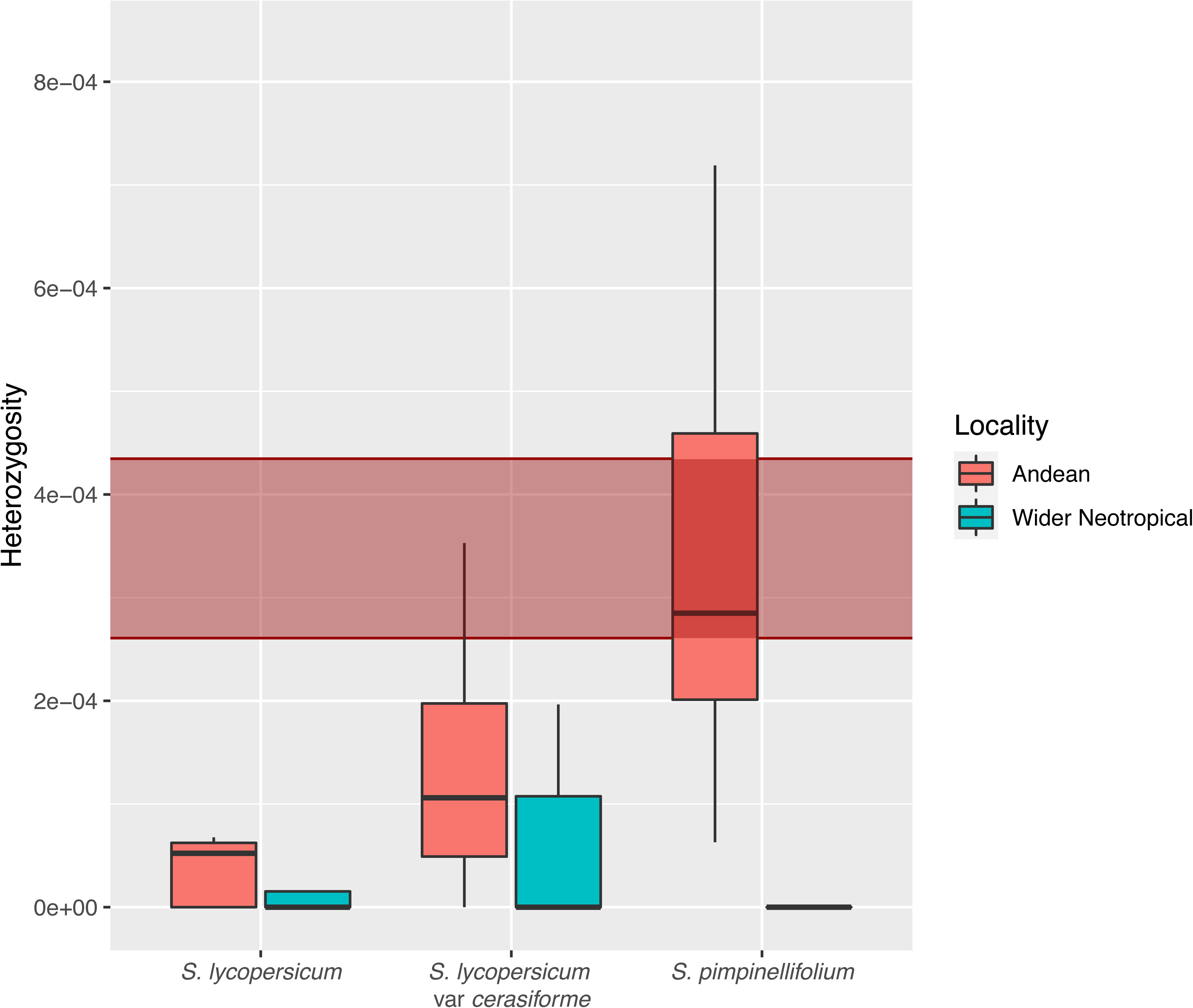
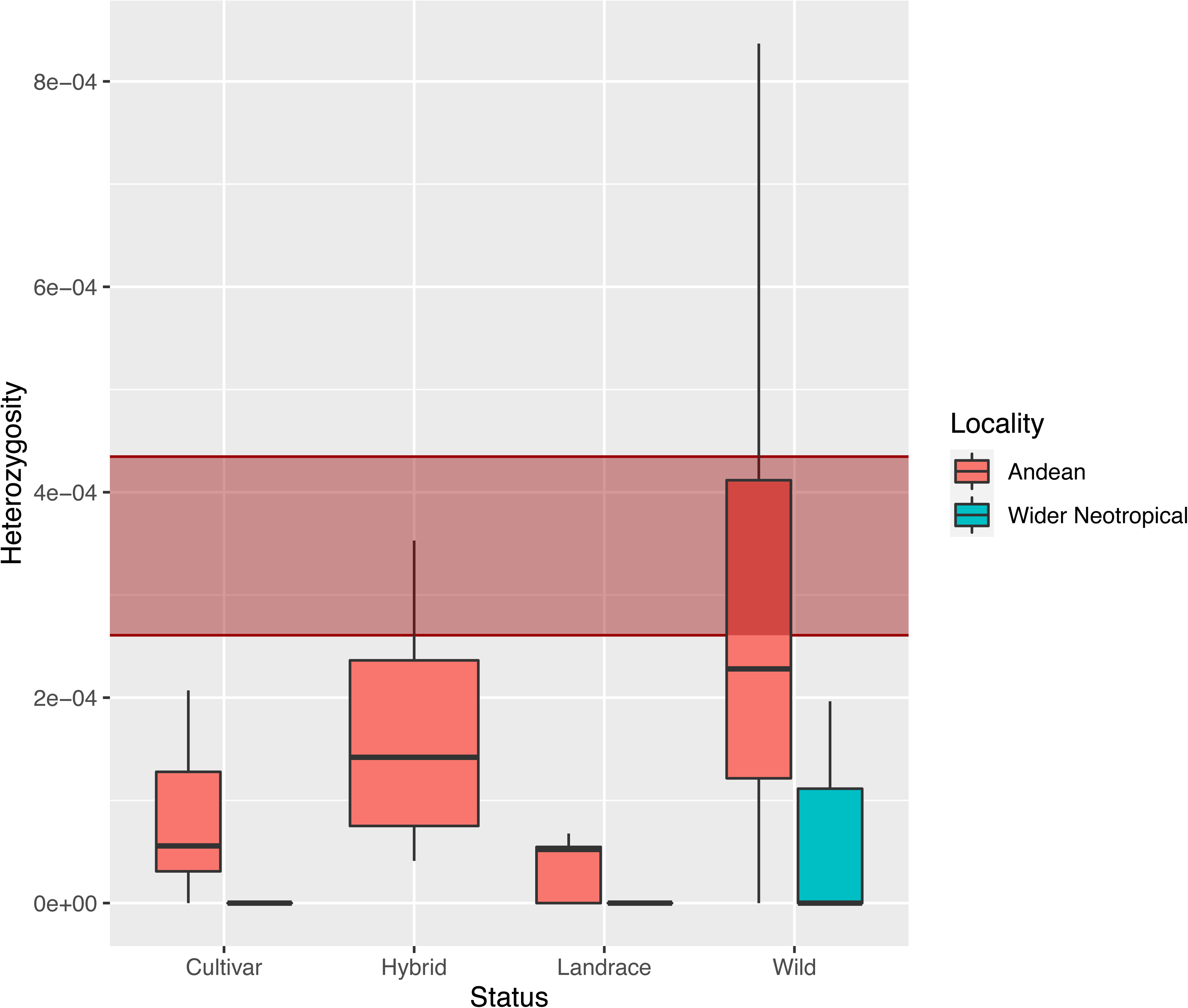
Box plots of heterozygosity partitioned by taxonomic variant (a) and botanical status (b). In both plots, each category is further partitioned by broad geographic origin. The shaded area represents the interval for the estimated heterozygosity in the *En Tibi* specimen.

## Discussion

The *En Tibi* tomato specimen is historically unique and of invaluable cultural and scientific significance (46, 47). Therefore, we minimised the amount of material we sampled destructively in our study. Combined with the organic decay processes and physical handling to which the specimen has been exposed for nearly a half millennium, this has resulted in an assembly with low coverage and the hallmarks of DNA fragmentation and modest post-mortem modification. Ongoing improvements in ancient DNA lab protocols and high-throughput Next Generation Sequencing (NGS) will uncover more data in the future (48). We look forward to the advances in these techniques in this exciting time for archeogenomics.

Nevertheless, current NGS platforms yield good results. Due to the parallelism inherent in these platforms, decay signatures (fragmentation, post-mortem modification (31)) apparent in one template molecule are swamped in the assembly by observations from other molecules if sufficient numbers of these are available, resulting in inferences such as we present here. Our analyses incorporated multiple strategies to mitigate further the biases that decay processes might introduce. In the network analysis shown in Figure 3, we only present SNPs shared by multiple accessions, following the rationale that post-mortem modified bases in the *En Tibi* specimen would likely present as autapomorphies in our data. In the results shown in Figure 6, we include a conservative estimate of heterozygosity for the specimen resulting from a critical appraisal of the putatively heterozygous alleles, i.e. the numerator in the ratio of heterozygous over homozygous allele counts, discarding any with low coverage or at sites and with bases typical of post-mortem misincorporation.

The tantalising potential for reconstructing historical plant and fruit traits from functional genomics eluded us in this study. Although we have performed pilot studies on agronomically interesting genes and looked for GO term enrichment induced by coding SNPs in the *En Tibi* specimen (49), we omit these results here as they were highly inconclusive due to the low coverage both in terms of genomic regions covered at all, and the depth at which these were mapped. Because phylogenomic network construction, ancestral population structure analysis and calculations of heterozygosity can be performed on disparate, neutral loci, these were possible with the genome skimming approach available to us with the present data.

The results we have obtained derive from comparative analyses where the *En Tibi* specimen is placed in the context of deeply sequenced accessions enriched with metadata about sampling locality, cultivation, and botanical status. As resequencing projects proceed, greater resolution in some of our inferences may become possible. At present, we were somewhat hampered by the incompleteness of some of the metadata in the current panels. For example, many accessions lack georeferencing beyond sampling region or city. This meant that a more precise triangulation of the genetic origin of the *En Tibi* specimen analogous to results obtained in other research (21) was not possible.

In addition to the sparseness of some metadata, we also note that metadata semantics were sometimes ambiguous. For example, some specimens annotated as ‘wild’ tomatoes should be considered feral instead. What constitutes a ‘landrace’ as opposed to a ‘cultivar’ is likewise not clearly defined in our panel. Unfortunately, it may be impossible to remedy this.

Many of the accessions in our panel are obtained from historical gene bank material curated by the C. M. Rick Tomato Genetics Resource Center at UC Davis (USA). Often, these accessions were collected by Dr Charles M. Rick (1915-2002), in the field or from indigenous markets, during the middle decades of the twentieth century (50). The ongoing replacement of indigenous heirloom tomatoes by industrial cultivars, the destruction of natural and cultural habitats, and the loss of knowledge both indigenous and from early natural historians mean that some information available back then is lost forever, unrecoverable by further sampling and resequencing.

A further source of ambiguity in the metadata was the status of C tomatoes. The accessions in this group showed great variety in their placement in the network graphs (Fig. 3 and 4), with some showing affinity with P and others with B. The edge lengths were also diverse, with some accessions showing lengths resembling those of the P accessions (Fig. 3); heterozygosity in C was greater than in B (Fig. 6). Lastly, the fastSTRUCTURE analysis (Fig. 5) showed that, qualitatively, the C accessions were less characterised by the influence of a single ancestral population than B or P.

## Conclusions

Here we present the ancient nuclear and plastid genomes of a tomato specimen from the historic *En Tibi* herbarium dated to ca. 1558 (17, 18). We place these genomes within the context of the diversity of the wider tomato section *Lycopersicon* and the narrower complex of *Solanum pimpinellifolium* L., and the feral and cultivated varieties of *S. lycopersicum* L. (large-fruited tomatoes) and *S. lycopersicum* var *cerasiforme* L. (cherry tomatoes) that are derived from it. For the feral and cultivated varieties, we restrict ourselves to accessions from the Neotropics. Although the nuclear genome coverage was low and showed some of the hallmarks of post-mortem molecular decay processes, we can draw several conclusions.

The NeighborNet analysis resulted in the placement of the nuclear genome (Fig. 3) in accordance with what the phenotype of the specimen indicates, namely that this is a large-fruited tomato, not a cherry, like its nearest two neighbours. The genome is placed among three accessions collected around the Gulf of Mexico. The genetically closest one, B.153, is TGRC accession LA1544 collected in Campeche on the Yucatán Peninsula. The second nearest, B.249 (LA1462), was collected in Mérida, also on the Yucatán Peninsula. The third, C.233 (LA1218), originates from Vera Cruz on the Mexican mainland facing the Gulf. These results suggest that the *En Tibi* tomato derives from a variety that was cultivated in eastern Mexico, possibly the Yucatán Peninsula, whence germplasm was transported to Europe during the early Columbian Exchange (10).

The median-joining network analysis of the concatenated variable plastid markers (Fig. 4) is less resolved, thereby illustrating the large-scale trends in tomato domestication that other studies have recently uncovered. Here, the *En Tibi* specimen is placed in a cluster of low genetic diversity that comprises nearly all Mesoamerican accessions and numerous South American ones. The pattern of low genetic diversity with high geographic dispersal is characteristic of domestication. This cluster thus represents the later stage of Mesoamerican tomato domestication that multiple studies posit, either preceded by earlier Andean predomestication (5) or the natural evolution of *S. lycopersicum* var. *cerasiforme* in the Andes (6).

The median-joining network also shows a central cluster of Peruvian, Ecuadorian and Bolivian accessions of *S. pimpinellifolium* L. and *S. lycopersicum* var. *cerasiforme* L. that is broadly compatible with both scenarios. However, the greater genetic diversity around the central node is notable and suggests processes that are substantially older and more lengthy than those that took place in Mesoamerica. The promiscuity among the members of the complex *S. pimpinellifolium* L., *S. lycopersicum* L. and *S. lycopersicum* var. *cerasiforme* L. that manifests as the interdigitation of the accessions in Figures 3 and 4 hampers drawing firm conclusions. Nevertheless, we note that some of the *cerasiforme* accessions have long branches and are placed among *S. pimpinellifolium* L., while others have short branches and occur among large-fruited cultivars. The former might be ‘true’ *cerasiforme* in the taxonomic sense (as per Razifard et al., (6)), while the latter might only be so in the phenetic sense, being merely small-fruited cultivars.

The fastSTRUCTURE analysis (Fig. 5) further illustrates the promiscuity within the complex, as the three hypothesised ancestral populations are components that shape all three members of the complex. Nevertheless, the geographic structure (Fig. 5b) of the ancestry components in *S. pimpinellifolium* L. is interesting. The southern population is underrepresented in domesticated tomatoes (Fig. 5b), suggesting that the onset of the C/B diversification history was in the northern Andean region. The reconstruction of an ancestral component within *cerasiforme* that is nearly absent from the large-fruited accessions is compatible with a natural origin of *cerasiforme* followed by domestication founder events that missed some of this diversity. The predominance of a third ancestral population in the ancestry of large-fruited tomatoes with a geographic centre south of the Gulf of Guayaquil invites hypotheses about human-assisted coastal dispersal as part of the second stage of tomato domestication that further, multidisciplinary research may elucidate.

The heterozygosity analysis indicates that the *En Tibi* specimen is highly heterozygous, and this is possibly underestimated given the conservative estimate of heterozygous site counts we generated to mitigate the effects of post-mortem decay processes. Similar considerations played into our approach to the NeighborNet construction, where we included only shared states. In pilot network analyses where we also included autapomorphies, the terminal branch leading to the *En Tibi* was notably longer than that of near neighbours (results not shown). Determining the precise extent to which this results from post-mortem misincorporation versus genuine, unique alleles in the specimen is not straightforward. Nevertheless, the snapshot that the *En Tibi* provides of the dynamics of genetic diversity of domesticates through time offers no evidence for an hourglass model where diversity is reduced subsequent to domestication, followed by increasing diversity in concert with the expansion of cultivation and breeding. On the contrary, the genome is more compatible with Allaby et al.’s alternative model of monotonously decreasing diversity as time and cultivation proceeds (22). However, the heterozygosity in the *En Tibi* specimen may have come about in multiple ways. For example, the specimen may originate from a heterozygous, outbred population or, alternatively, from recent outcrossing of distinct, inbred cultivars. With greater contiguous coverage, these scenarios could be distinguished by the size of identical-by-descent segments. Unfortunately, with 0.75% of the reference genome covered across disparate sites, such analysis is not feasible at this time.

The genomic data of the *En Tibi* tomato specimen provides a snapshot of the dynamics of tomato domestication and cultivation through time and space, from the Andes and Mesoamerica in pre-Columbian times through the Columbian Exchange to Renaissance Europe. The insights that the specimen provides are only possible thanks to data and metadata provided by the specimen itself and other accessions with which it can be compared. The key specimens for this context are the result of Neotropical indigenous cultural and agricultural processes combined with collection, annotation and curation in gene banks and natural history collections. Absent all these ongoing contributions, the replacement of indigenous heirlooms by inbred industrial cultivars and the destruction of natural and cultural habitats in the Neotropics will erase the origins of one of the most important crops humankind has to offer.

## Supplementary Materials

The raw sequencing data generated in this study are deposited at the NCBI SRA as BioProject PRJNA566320. All other intermediate and result data are deposited in a collection of archives under DOI 10.5281/zenodo.4966843. Where applicable, we specify in the manuscript body which result set is contained in which archive. The source code developed to analyse the nuclear genome, principally by traversing the relational SNP database, is deposited under DOI 10.5281/zenodo.5940589. The source code for the chloroplast genome analysis is deposited under DOI 10.5281/zenodo.5940631.

## Acknowledgements

This work has been supported by in-kind contributions from Naturalis Biodiversity Center, Leiden, the Netherlands and Plant Research International at Wageningen University and Research, the Netherlands. We are grateful to the leadership of these respective institutions for facilitating this collaboration. The authors would like to acknowledge the editor and anonymous reviewers for their time and feedback on this manuscript.

