## Supplementary figures and images for "A Tomato Genome From The Italian Renaissance Provides Insights Into Columbian and Pre-Columbian Exchange Links And Domestication"

### Supplemental Figure 1

#### run0220_En−Tibi_S2_L003.fixmate.sorted.markdup

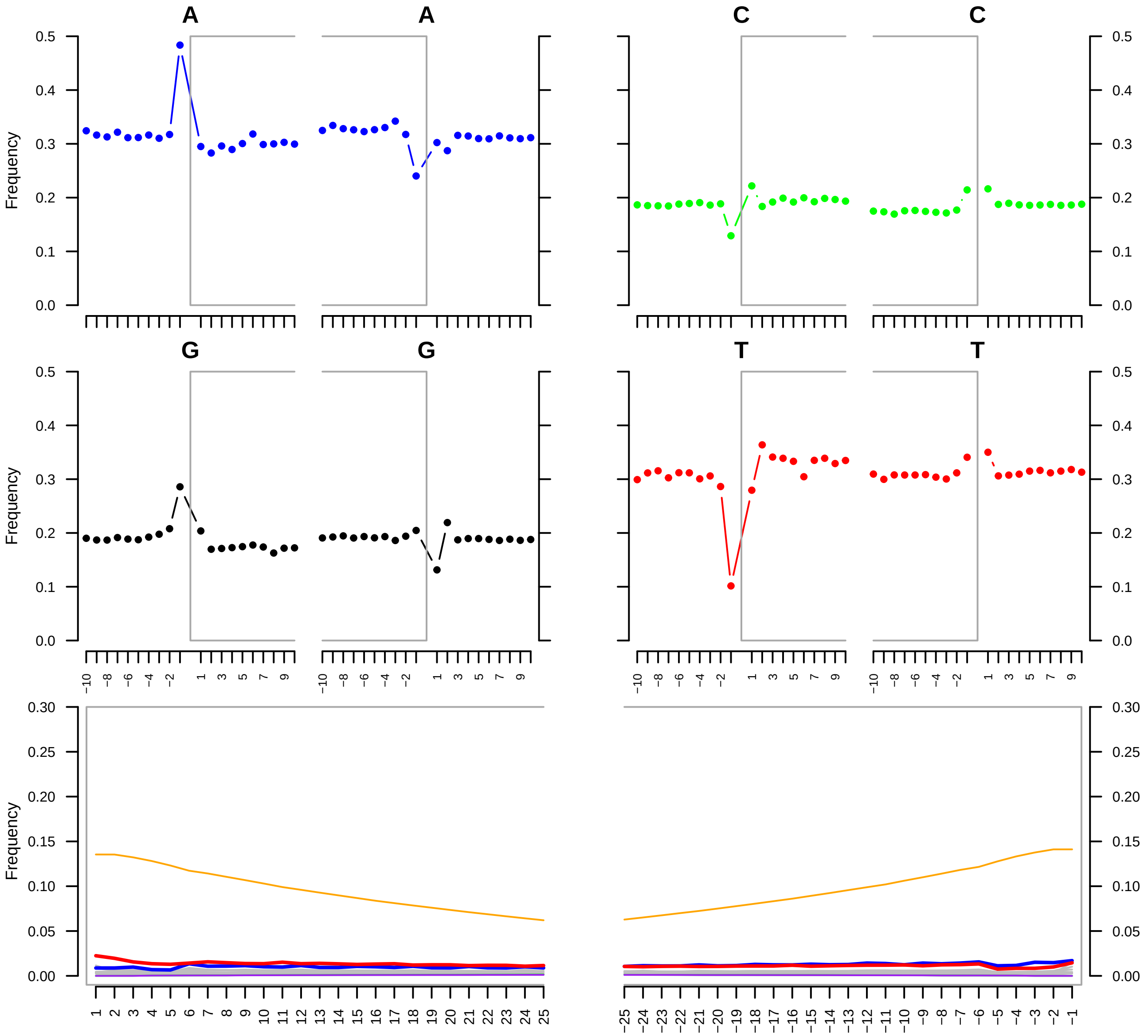
